# *In Vitro* Efficacy of Five Biofungicides against *Rhizoctonia solani* Kuhn in Rice

**DOI:** 10.1101/2020.07.22.216325

**Authors:** Irish Mae Bauzon-Cantila, Jaime C. Silvestre, Raquel B. Evangelista, Edralyn Catubay

## Abstract

*Rhizoctonia solani* Kuhn, the causal pathogen of sheath blight is second most damaging fungal disease in rice. While using chemical fungicides present high detriment to environment, the study investigate the efficacy of treatments composing five biofungicides in three different rates along with a biological agent, chemical check and untreated against the pathogen in *in vitro* level. *In vitro* efficacy showed that *Melaleuca alternifolia* + terpenes at 3.00 ml/L of H_2_0 (T6), *Aloe vera* powder (Manopol) + *Melaleuca* oil at 3.00 ml/L of H_2_0 (T15) and at 2.00 ml/L of H_2_0 (T14) and *Melaleuca alternifolia* + terpenes at 2.00 ml/L of H_2_0 (T5) as very effective (0-10 mm diameter zone of growth) treatments comparable to the chemical check (T17). Therefore, attaining high yield rice while having low risk to environment can always be done.

## INTRODUCTION

*Rhizoctonia solani* Kuhn, causing sheath blight is one of the most economically important fungal diseases that limits rice production [1], which its sclerotia or hyphae can easily attach to the rice sheath and infecting blight in rice [2]. Further infections are caused by hyphae growing upward towards uninfected plant parts and produce additional lesions and sclerotia on leaf sheaths affecting productivity in plants [3, 4]. Depending on the plant stage upon infection and type of environment, a range of 4 to 50% can be reduced to the yield, making sheath blight as the second most damaging fungal disease in rice [1, 5-8].

Rice released varieties such as hybrid rice and foreign germplasm with low resistance to sheath blight [9] had been tested to high rainfall areas in the Philippines [10, 11], which is a disease prone environment. It could be advantageous then to apply biofungicides to the crop. Biofungicides as one of the methods that can manage the disease is gaining popularity because it is more environment-friendly [12, 13]. Moreover, regulations towards utilizing chemical fungicides could be stricter in the future as several organizations are putting pressure on taking out hazardous chemicals in the market [13]. Hence, this study was conducted to investigate the *in vitro* efficacy of biofungicides against *R. solani* Kuhn in rice.

## MATERIALS AND METHODS

### INOCULUM AND TREATMENT PREPARATION

Diseased sheaths of rice due to blight were collected in the field (Figure 1) and placed into plastic bag for diagnosis and isolation. Potato dextrose agar (PDA) was then used as medium for isolating sheath blight pathogen with the following components: 200 g of clean potato, 1 L of water, 20 g dextrose, and 20 g agar. The medium was dispensed in flat bottles, plugged with cotton and autoclaved at 15 *psi* or 121°C for 15-30 minutes. Tissue planting technique was used to transfer pathogen in PDA medium, which was incubated under room condition. Purification was done by transferring mycelial bits onto fresh PDA medium.

**Figure 1.**
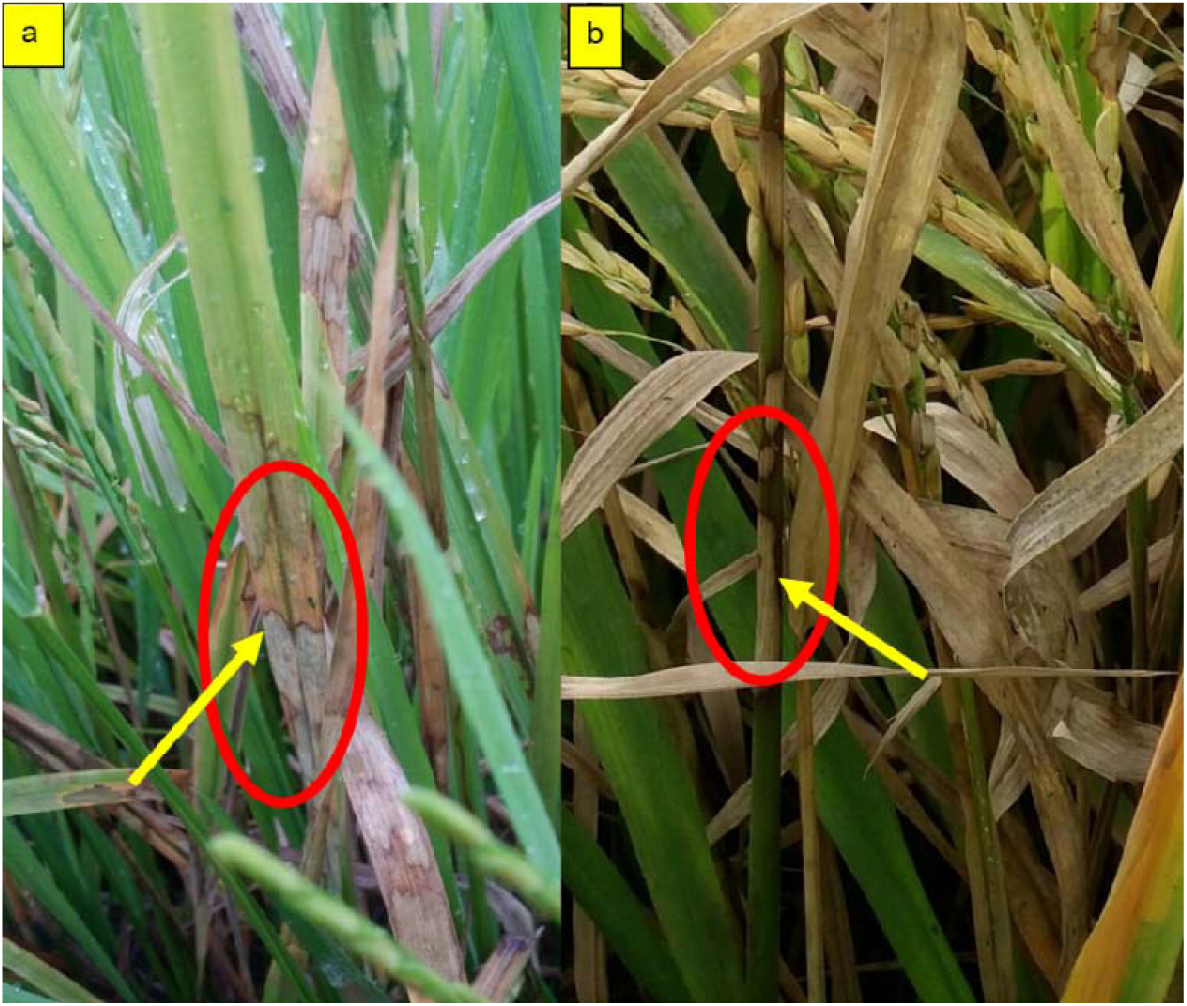
Rice plants showing symptoms of sheath blight on the (a) leaf (gray irregular lesions banded with green brown coloration and grayish center) and (b) sheath (oval gray spots with black brown margins that later enlarge).

A total of 18 treatments composed of five biofungicides in three different rates, a biological control, chemical check, and untreated was used. The main component of the five biofungicides is *Bacillus subtilis, Melaleuca alternifolia* + terpenes, seaweed extract, *Lactobacillus plantarum*, aloe vera powder (Manopol) + *Melaleuca* oil. *Trichoderma* spp. was used as biological agent while Propiconazole + difenoconazole as chemical check. Other details of each treatment can be found in Table 1.

**Table 1.** Biofungicides and their corresponding details that has been used in the study.

| Component | Commercial name | Reports | Source | Treatment (rate*) |
| --- | --- | --- | --- | --- |
| *Bacillus subtilis* | Seranade 250 | suppresses club root to canola on *in vitro* conditions and effective against crop damaging pathogens like powdery mildew, walnut blight and *Botrytis* bunch rot and fire blight | [18, 19] | T1 (1.25 ml) |
|  |  |  |  | T2 (2.5 ml) |
|  |  |  |  | T3 (3.75 ml) |
| *Melaleuca alternifolia* + terpenes | Timorex Gold | having prophylactic and curative activity against broad spectrum of diseases in crops | [20] | T4 (1 ml) |
|  |  |  |  | T5 (2 ml) |
|  |  |  |  | T6 (3 ml) |
| Seaweed extract | MPQ | produces metabolites with antiviral, antiprotozoal, antifungal, and antibacterial properties aiding in the protection | [21, 22] | T7 (0.25 g) |
|  |  |  |  | T8 (0.5 g) |
|  |  |  |  | T9 (0.75 g) |
| *Lactobacillus plantarum* | EM-1 | produce antifungal substances such as benzoic acid, methylhydantoin, mevalonolactone against plant pathogens | [23-25] | T10 (1 ml) |
|  |  |  |  | T11 (2 ml) |
|  |  |  |  | T12 (3 ml) |
| *Aloe vera* powder (Manopol) + *Melaleuca* oil | AZ 41 | effectively control diseases and insect pests in citrus | [17] | T13 (1 ml) |
|  |  |  |  | T14 (2 ml) |
|  |  |  |  | T15 (3 ml) |
| Biological control | *Trichoderma* spp. | induce systemic resistance against plant pathogens | [26] | T16 (0.5 ml) |
| Propiconazole + difenoconazole | Armure | controls *Rhizoctonia solani* Kuhn in rice very effectively | [27, 28] | T17 (1 ml) |
| Untreated |  |  |  | T18 (Untreated) |

\*diluted in L of H<sub>2</sub>O, MPQ= Marvelous Premium Quality, EM= effective microorganism

### BIOASSAY TEST

In a 14 days old *R. solani* Kuhn culture (Figure 2), culture disc method was used for fungicidal assay with the following steps in Figure 3. Petri plates were rotated thoroughly for mixing and congealing. Culture disc of the pathogen was placed at the center of the plates, which were labelled and incubated in inverted position under room temperature. Measurement of the diameter zone of growth (mm) of the pathogen was done after 72 hrs of incubation.

**Figure 2.**
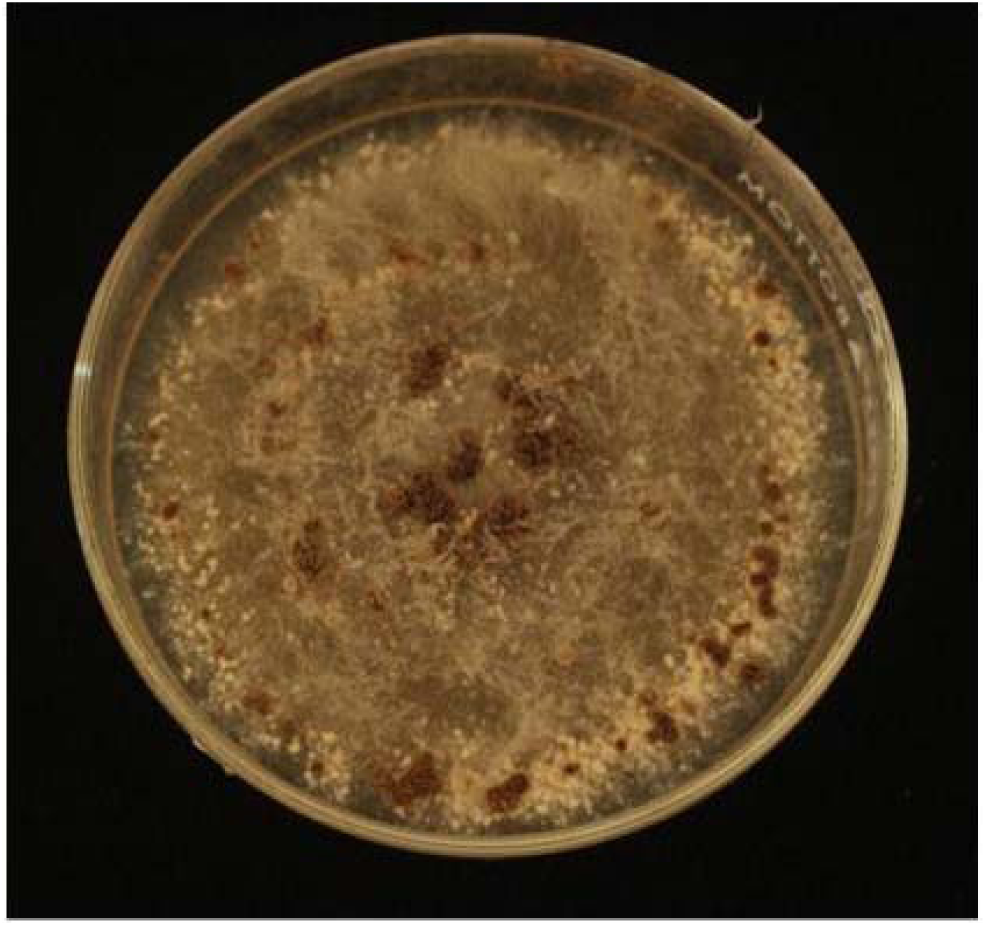
Pure culture of *Rhizoctonia solani* Kuhn in 14 days old showing mycelia and matured sclerotia on potato dextrose agar.

**Figure 3.**
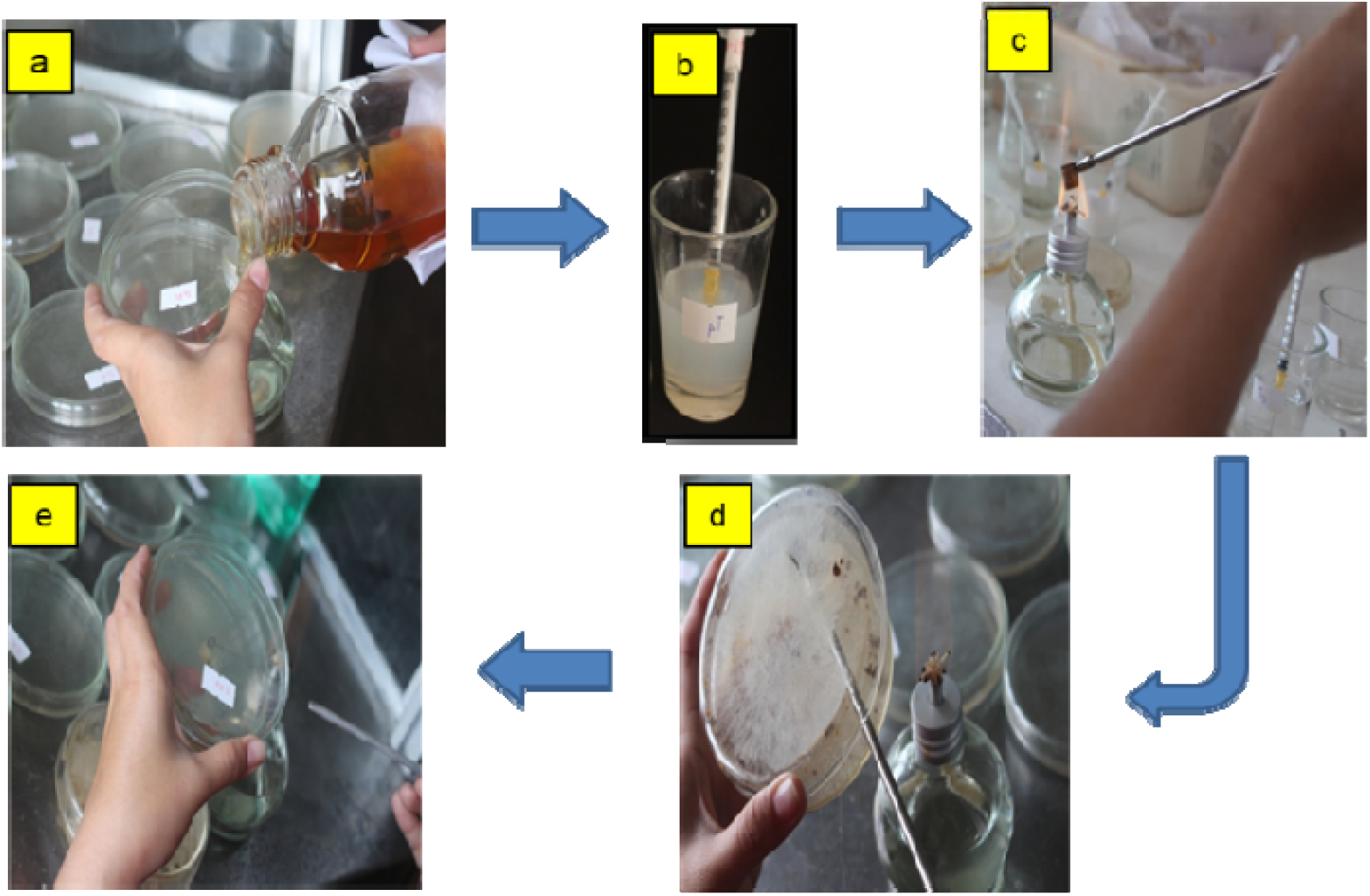
Culture disc method for biofungicidal assay of *Rhizoctonia solani* Kuhn. *a) Potato dextrose agar (PDA) was used as medium for isolating the pathogen, b) 1 ml of the treatment was added to each petri plate with 20 ml melted PDA, c) Ensure aseptic condition by flaming the corkborer, d) The isolated pathogen was removed aseptically, and e) The culture disc of the pathogen was placed at the center of each plate.

### DATA ANALYSIS

The *in vitro* test was laid-out in completely randomized design with 18 treatments in triplicates. The data gathered was diameter zone of growth (DZG), which is a measurement to determine the efficacy of treatment. The data (mm) was taken after 72 hrs of incubation, which efficacy follows an arbitrary scale range: 0-10 mm is very effective, 11-20 mm is effective, 21-30 mm is moderately effective, and 31 mm and above is not effective. The analysis of variance and mean separation were done using Statistix 9.0 [14].

## RESULTS AND DISCUSSION

*In vitro* result showed the mean DZG (mm) of *R. solani* Kuhn was significantly affected by the application of different rates of five biofungicides after 72 hrs of incubation (Table 2). The most effective biofungicide was *Melaleuca alternifolia* + terpenes at 3.00 ml/L of H_2_0 (T6) with no observed growth of the pathogen (Figure 4). *Aloe vera* powder (Manopol) + *Melaleuca* oil at 3.00 ml/L of H_2_0 (T15) with 0.17 mm and at 2.00 ml/L of H_2_0 (T14) with 2.17 mm and *Melaleuca alternifolia* + terpenes at 2.00 ml/L of H_2_0 (T5) with 3.83 mm followed as among the most effective treatments (Figure 4). The effect of *Melaleuca alternifolia* + terpenes (T5 and T6) and *Aloe vera* powder (Manopol) + *Melaleuca* oil (T14 and T15) were comparable to the chemical check (T17) and rated as very effective (Figure 4). This implies that these two biofungicides were as effective as the chemical check in inhibiting the growth of *R. solani* Kuhn *in vitro*.

**Table 2.** Diameter zone of growth (mm) of *R. solani* Kuhn as influenced by different rates of biofungicides

| Treatments | Mean (mm) <sup>1</sup> | Degree of Effectiveness |
| --- | --- | --- |
| T1 | 18.17 <sup>cde</sup> | Effective |
| T2 | 20.50 <sup>bcde</sup> | Moderately effective |
| T3 | 22.17 <sup>bcde</sup> | Moderately effective |
| T4 | 14.83 <sup>de</sup> | Effective |
| T5 | 3.83 <sup>e</sup> | Very effective |
| T6 | 0.00 <sup>e</sup> | Very effective |
| T7 | 53.17 <sup>ab</sup> | Not effective |
| T8 | 66.83 <sup>a</sup> | Not effective |
| T9 | 48.83 <sup>abcd</sup> | Not effective |
| T10 | 44.00 <sup>abcd</sup> | Not effective |
| T11 | 40.83 <sup>abcd</sup> | Not effective |
| T12 | 41.33 <sup>abcd</sup> | Not effective |
| T13 | 20.33 <sup>bcde</sup> | Effective |
| T14 | 2.17 <sup>e</sup> | Very effective |
| T15 | 0.17 <sup>e</sup> | Very effective |
| T16 | 51.67 <sup>abc</sup> | Not effective |
| T17 | 0.00 <sup>e</sup> | Very effective |
| T18 | 68.33 <sup>a</sup> | Not effective |

<sup>1</sup>Means in column followed by common letter superscripts are not significantly different at 1% level, Tukey's Test.

**Figure 4.**
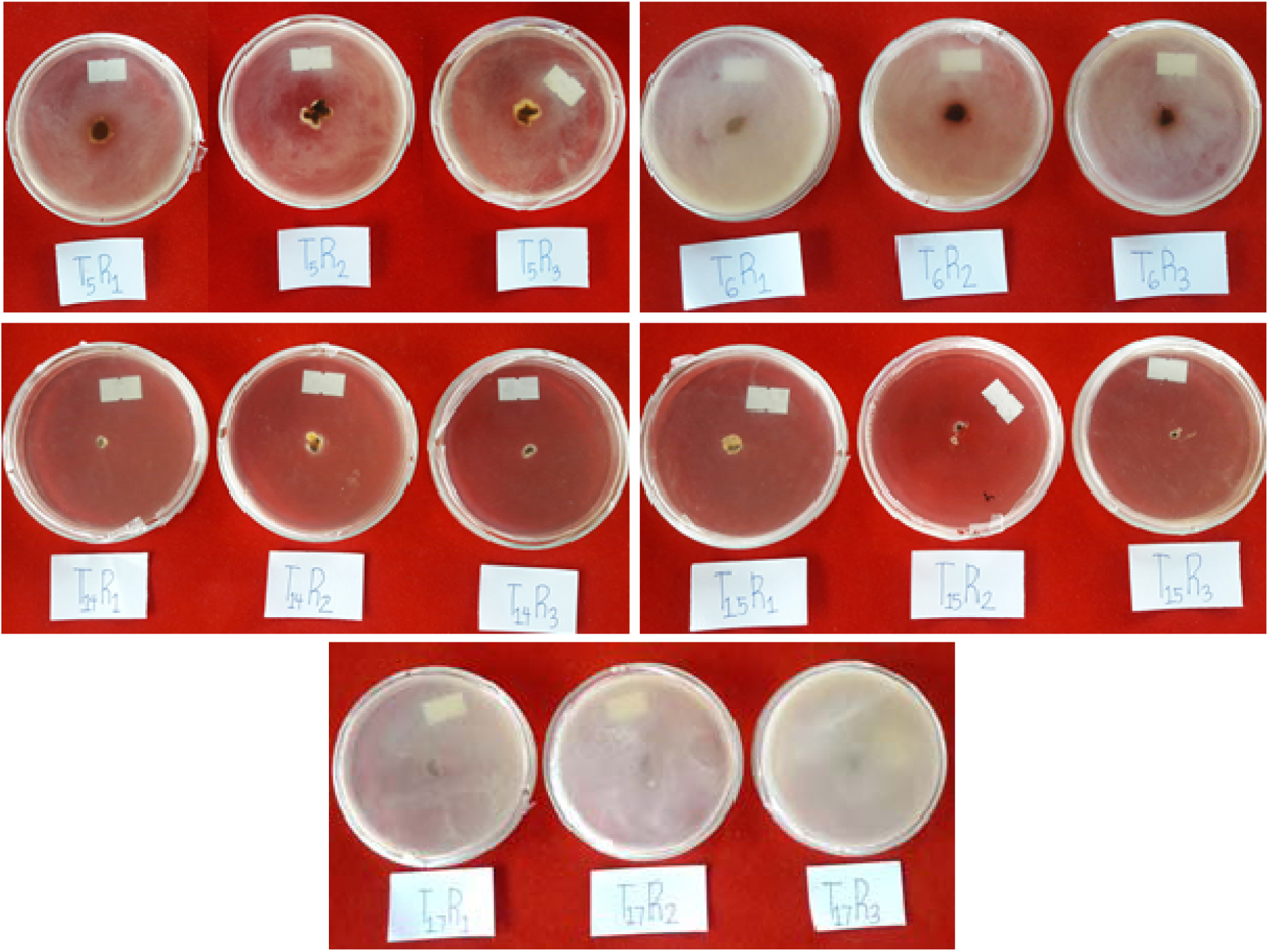
Diameter zone of growth of *R. solani* Kuhn showing very effective treatments in petri dish.

Previous reports showed that *Aloe vera* powder (Manopol) + *Melaleuca* oil effectively controlled *Corynespora* leaf fall disease of rubber caused by *Corynespora cassiicola* in *in vitro* study [15] while *Melaleuca alternifolia* + terpenes was found effective against *Pythium aphanidermatum* on pepper [16] and effectively control diseases *Citrus madurensis* [17]. The lowest rates at 1.00 ml/L of H_2_0 of *Melaleuca alternifolia* + terpenes (T4) and *Aloe vera* powder (Manopol) + *Melaleuca* oil (T13) were still rated effective, similar to the effect of lowest rate of *Bacillus subtilis* at 1.25 ml/L of H_2_0 (T1) (Figure 5). *Bacillus subtilis* at 2.5 ml/L of H_2_0 (T2) and 3.75 ml/L of H_2_0 (T3) were rated as moderately effective (Figure 6). Lastly, the three rates of seaweed extract and *Lactobacillus plantarum* were rated not effective with DZG means ranging from 40.83 to 66.83 mm and their results were comparable to the untreated control and biological check with DZG means of 68.33 and 51.67 mm, respectively (Figure 7).

**Figure 5.**
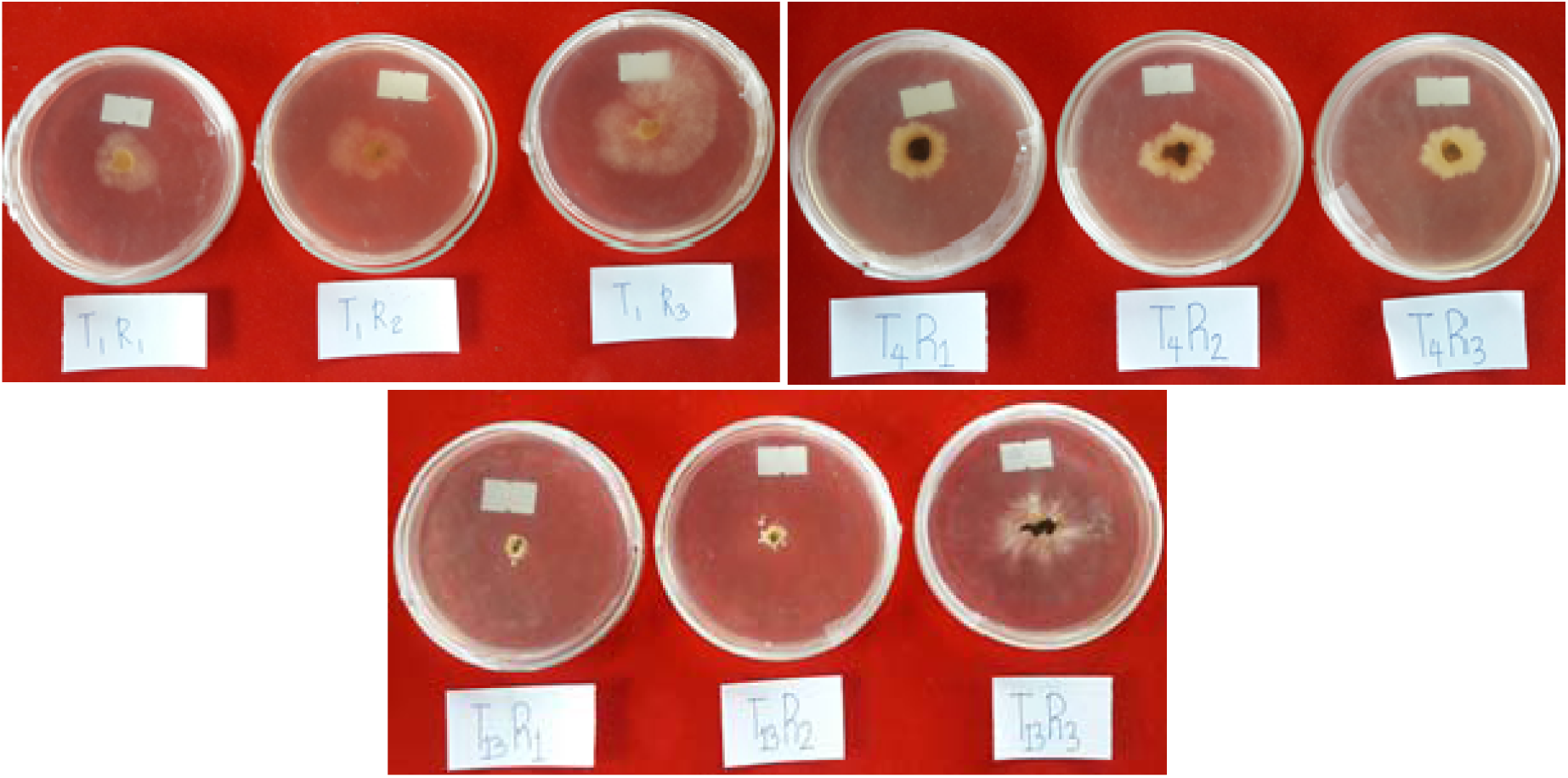
Diameter zone of growth of *R. solani* Kuhn showing effective treatments in petri dish.

**Figure 6.**
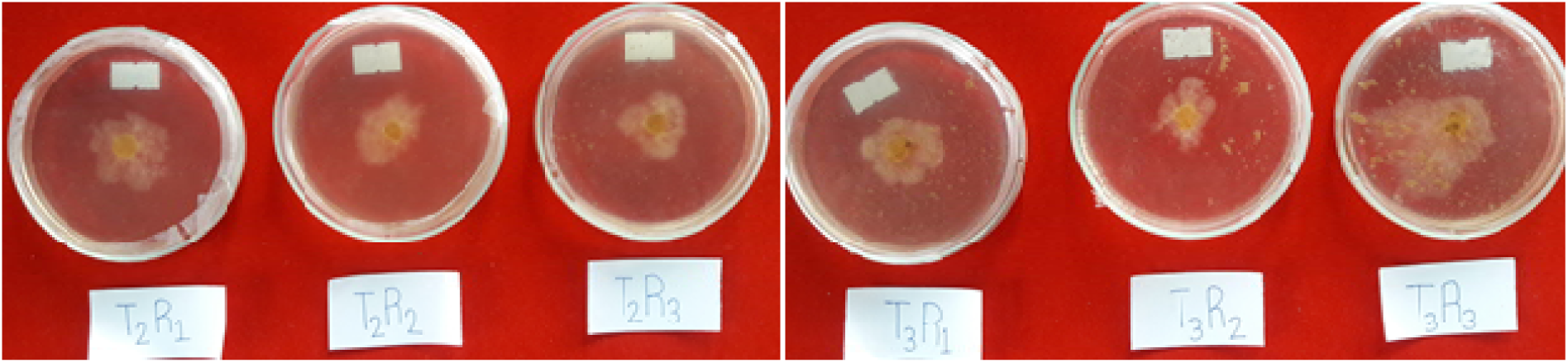
Diameter zone of growth of *R. solani* Kuhn showing moderately effective treatments.

**Figure 7.**
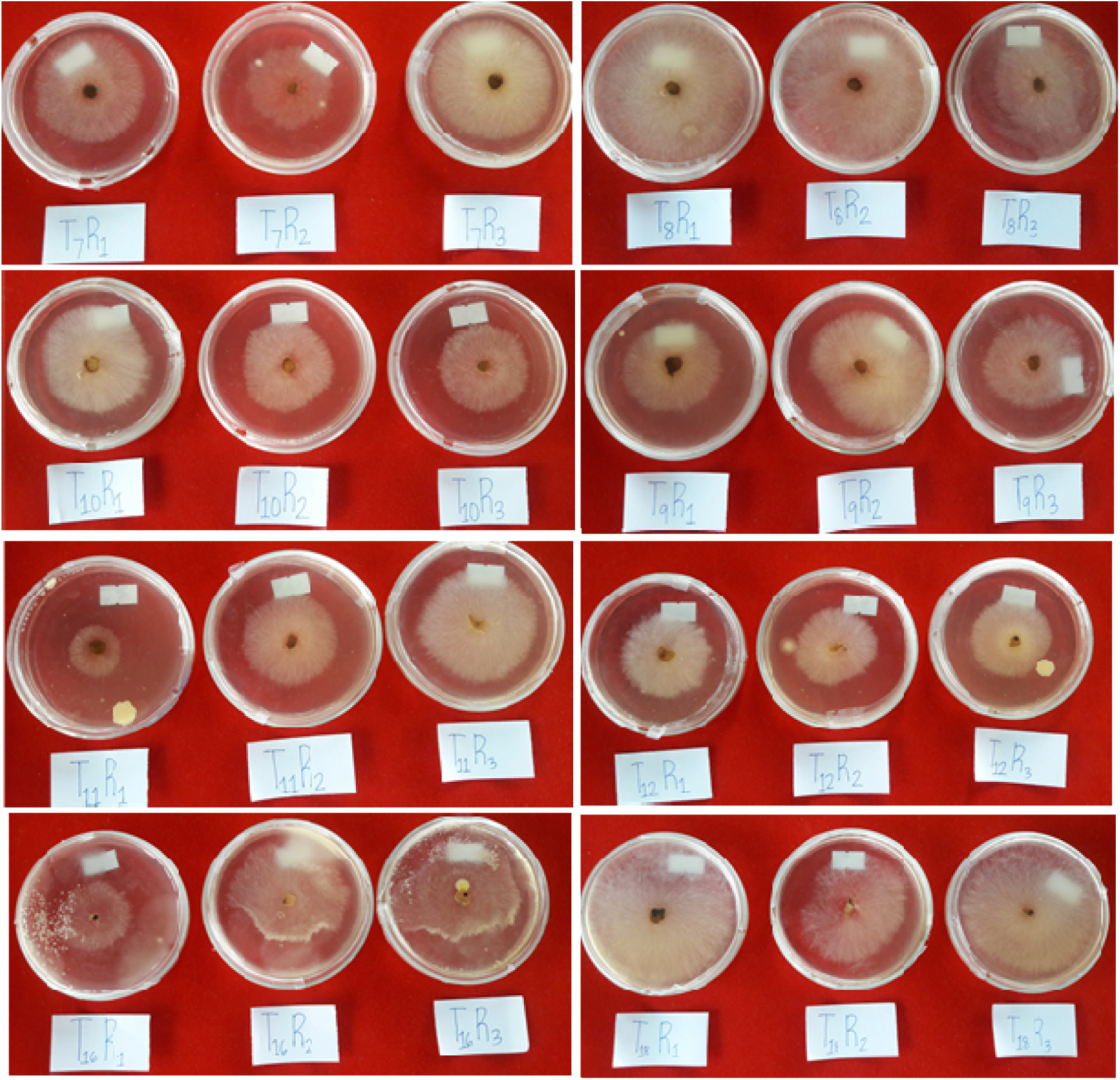
Diameter zone of growth of *R. solani* Kuhn showing not effective treatments.

## CONCLUSION

Biofungicides especially those containing *Melaleuca alternifolia* and *Aloe vera* are very effective in controlling *R. solani* Kuhn in rice in *in vitro*, which could be verified in *in vivo* study. Attaining high yield and less pesticides residue in rice is always possible, which should be promoted to uphold a safer environment.

## ACKNOWLEDGEMENTS

The authors would like to thank Department of Plant Pathology, College of Agriculture for allowing the utility of laboratory equipment.

## STATEMENT ON CONFLICT OF INTEREST

The authors declare no conflict of interest.

## Notes

### Competing Interest Statement

The authors have declared no competing interest.

